# Temporal Effects of Galactose and Manganese Supplementation on Monoclonal Antibody N-Linked Glycosylation in Fed-Batch and Perfusion Bioreactor Operation

**DOI:** 10.1101/2023.04.15.535602

**Authors:** Aron Gyorgypal, Erica Fratz-Berilla, Casey Kohnhorst, David N. Powers, Shishir P. S. Chundawat

**Author notes:** **Corresponding Author:** Shishir P. S. Chundawat.

## Abstract

Monoclonal antibodies (mAbs) represent a majority of biotherapeutics on the market today. These glycoproteins undergo post-translational modifications, such as N-linked glycosylation, that influence the structural & functional characteristics of the antibody. Glycosylation is a heterogenous post-translational modification that may influence therapeutic glycoprotein stability and clinical efficacy, which is why it is often considered a critical quality attribute (CQA) of the mAb product. While much is known about the glycosylation pathways of Chinese Hamster Ovary (CHO) cells and how cell culture chemical modifiers may influence the N-glycosylation profile of the final product, this knowledge is often based on the final cumulative glycan profile at the end of the batch process. Building a temporal understanding of N-glycosylation and how mAb glycoform composition responds to real-time changes in the biomanufacturing process will help build integrated process models that may allow for glycosylation control to produce a more homogenous product. Here, we look at the effect of specific nutrient feed media additives (e.g., galactose, manganese) and feeding times on the N-glycosylation pathway to modulate N-glycosylation of a Herceptin biosimilar mAb (i.e., Trastuzumab). We deploy the N-GLYcanyzer process analytical technology (PAT) to monitor glycoforms in near real-time for bench-scale bioprocesses operated in both fed-batch and perfusion modes to build an understanding of how temporal changes in mAb N-glycosylation are dependent on specific media additives. We find that Trastuzumab terminal galactosylation is sensitive to media feeding times and intracellular nucleotide sugar pools. Temporal analysis reveals an increased desirable production of single and double galactose-occupied glycoforms over time under glucose-starved fed-batch cultures. Comparable galactosylation profiles were also observed between fed-batch (nutrient-limited) and perfusion (non-nutrient-limited) bioprocess conditions. In summary, our results demonstrate the utility of real-time monitoring of mAb glycoforms and feeding critical cell culture nutrients under fed-batch and perfusion bioprocessing conditions to produce higher-quality biologics.

## Introduction

Monoclonal antibodies (mAbs) produced from Chinese hamster ovary (CHO) cells represent a majority of biotherapeutics on the market today and continually exhibit steady annual growth rates. In addition to the approval of new innovators entering the market every year, there has been a rise within the biosimilar market as patent protections expire for older mAb products.^1,2^ The ability of CHO cells to produce humanized or human-like post-translational modifications, such as N-linked glycosylation, makes them highly desirable as an expression system for producing mAb products.

N-linked glycosylation is a common post-translational modification where specific oligosaccharides (i.e., N-glycans) are attached to an asparagine residue on the protein backbone of the Fc region of the mAb. The N-glycans attached to antibodies are known to influence their clinical efficacy, such as stability, pharmacodynamics, and pharmacokinetics.^3,4^ The biosynthesis of glycoproteins in mammalian cells involves a complex and interconnected network of glycosylation enzymes within the secretory compartments of the endoplasmic reticulum (ER) and the Golgi apparatus. The glycosylation pathway is also sensitive to cellular metabolism, thus producing heterogeneous glycosylation of the secreted mAb product. Perturbations during bioprocessing (i.e., changes in the dissolved oxygen (DO) levels, pH, temperature, and agitation rates) can influence the pathways as well, influencing the final glycan profile, often making it important to identify the critical process parameters (CPP) and critical material attributes (CMA) during upstream bioprocessing that can influence final drug CQAs like N-glycosylation.^5–9^

The N-glycans attached to a mAb protein can often be characterized or differentiated by its terminal sugars, which influence final drug clinical characteristics. For example, afucosylated mAbs have been reported to have improved binding to the FcγRIII receptor as a consequence of the absence of the molecule, increasing accessibility for interaction and thus increasing Fc-mediated antibody-dependent cellular cytotoxicity (ADCC).^10,11^ Terminal galactose branching enhances complement-dependent cytotoxicity (CDC) by increasing its binding activity to C1q over its non-galactosylated counterparts.^12^ Terminal sialic acids have been demonstrated to increase serum persistence and modulate anti-inflammatory activity.^13,14^ High mannose isoforms can potentially bind to serum mannose-binding lectin 2 and activate the lectin complement pathway on mAb-targeted cells, increasing their clearance *in-vivo*.^15,16^ Because the varying glycoforms can significantly impact the clinical efficacy of the mAb product, N-linked glycosylation is typically considered a CQA and needs to be closely monitored during bioprocessing.

Glycosylation profiles of the secreted mAb product can be altered using nutrient media feed additives that influence specific metabolic pathways inside CHO cells. Modulating galactosylation profile of mAbs using feed additives that impact glycan precursor levels in cells has been previously studied.^17–22^ Galactose (gal) and uridine are precursor molecules to produce Uridine diphosphate galactose (UDP-gal), the substrate for galactosyltransferase enzyme involved in the biosynthesis pathway. Manganese is a limiting cofactor necessary for this enzymatic reaction as well. Gramer et al. studied the effects of feeding uridine, manganese, and galactose at different bolus concentrations and observed a dose-dependent increase in the rate of mAb galactosylation that eventually reached saturation.^21^ A similar study by Kildegaard et al. observed that a 20 mM bolus galactose feed alone was enough to have a statistically significant impact on mAb galactosylation.^22^ Lastly, Sha et al. found that a galactose feed in lieu of glucose can increase the galactosylation rate of the mAb, which is influenced by an increase in the nucleotide sugar precursor UDP-gal concentrations within the cells.^19^ Other studies have also examined the influence of sialylation as a function of increased galactosylation^17^ or a combination of galactose with lactate on cell metabolism in relationship to mAb glycosylation. However, the influence of increasing terminal galactose glycan species did not increase the rate of sialylation as sialyltransferases often become a bottleneck in further modification of N-glycans.^18^

Other chemical modulators that do not directly feed into the glycosylation pathway have also been studied to understand their effects on mAb glycosylation.^23^ Sodium butyrate, a histone deacetylase inhibitor known to increase cell-specific monoclonal antibody production, has been widely studied, and an inverse relationship was found between cell-specific productivity and glycan maturation.^20^ More recently, rosmarinic acid has also been studied and shown to increase mAb titer favorably; however, no literature shows its effect on glycosylation.^24,25^

The temporal influence of nutrient feed addition during the cell culture process on the N-glycosylation process is still not fully understood for both fed-batch and continuous perfusion processes. This knowledge gap can be attributed to the lack of integrated process analytical technology (PAT) to allow real-time N-glycosylation analysis to monitor this critical quality attribute (CQA).^26,27^ We have recently showcased the N-GLYcanyzer PAT system that allows for near-real-time N-glycosylation analysis to fill this gap in process knowledge. Here, we look to first study the influence of the feeding regimen and the addition of galactose and manganese feed additives on the CHO cell process and N-glycosylation profile for a model mAb (Trastuzumab). Our experimental design incorporated fed-batch and perfusion bioreactor-based cell cultures with or without galactose/manganese supplementation to explore the impact of galactose availability on mAb glycosylation profiles (**Scheme 1**). We look to build an understanding of systematic glucose depletion and daily or alternative day intermittent refeeding with glucose/galactose/Mn nutrients in fed-batch mode operation and its effect on mAb N-glycosylation. With this knowledge, we next used the N-GLYcanyzer system during perfusion-mode bioreactor operation to also study the temporal changes in N-glycosylation profiles for the monoclonal antibody produced and its response to continuous glucose/galactose/Mn nutrient feeding.

## Materials and Methods

### Cell Line and Pre-inoculum

The Chinese hamster ovary (CHO-K1) cell line producing a recombinant mAb biosimilar of Trastuzumab was kindly donated by GenScript Biotech Corporation (Piscataway, NJ). A seed train was started by thawing one ampule of cells (1×10^7^ cells/mL) from the working seed bank into high-intensity perfusion CHO (HIP-CHO) medium (Thermo Fischer Scientific, Waltham, MA) containing 0.1% anticlumping agent (Thermo Fischer Scientific, Waltham, MA) in a 125 mL unbaffled shake flask (VWR, Radnor, PA, USA) with a 40 mL working volume. All bioreactor experiments were also conducted using this medium. The cells were grown at 37°C, 130 RPM, and 8% CO_2_ in a INFORS HT Multitron Incubator (Infors AG, Bottmingen, Switzerland) for 4 days and passaged twice to 0.5×10^6^ cells/mL into a 250 mL shake flask and then into a 500 mL shake flask, and then grown for 4 days before inoculation into the bioreactors for each experiment.

### Fed-Batch Bioreactor Culture

The fed-batch bioreactor culture experiments were run using an AMBR250 modular bioreactor (Sartorius Stedim North America, Bohemia, NY) with a starting volume of 200 mL. Temperature and pH control was set the day prior to inoculation to 37 °C and 7.1, respectively. The pH was controlled with bolus additions of 0.5 M NaOH and sparging of CO_2_. Dissolved oxygen (DO) was controlled at a setpoint of 50% using an O_2_ sparge as needed during the culture. The glucose or glucose/galactose feeds were prepared in HIP CHO Medium at a concentration of 250 g/L glucose or 250 g/L glucose and 75 g/L galactose. Feeding took place every 24 hours (daily) or 48 hours (alternative days) starting either day 3 or when the glucose concentration fell below 3 g/L, whichever came first. The galactose-supplemented cultures were also spiked to 1 μM manganese effective concentration using manganese chloride during the first feeding day to augment the glycosylation further, as manganese acts as a cofactor for galactosylation.

Cultures were fed to 5 g/L glucose effective concentration at specified intervals. A 3% (v/v) Antifoam C Emulsion solution (Sigma-Aldrich, St. Louis, MO) was added manually as needed. The bioreactors were inoculated to an initial density of 0.5 × 10^6^ cells/mL. Daily samples were taken to analyze various culture parameters (e.g., glucose, lactate, glutamate, glutamine, Na^+^, K^+^, and Ca^2+^) on a BioProfile FLEX2 Analyzer (Nova Biomedical, Waltham, MA). Galactose concentrations were measured using a colorimetric galactose assay kit (ka1669, Abnova, Taipei City, Taipei, Taiwan). The FLEX2 Analyzer was also tested for galactose interference when measuring glucose. Although galactose presence did increase the glucose reading at high concentrations (above 4.0 g/L galactose), it had no meaningful impact at the lower concentrations under 2 g/L. As our study looked at galactose concentrations at or below 1.8 g/L, there was no meaningful skew in the glucose presence for these studies.

Titer analysis of spent medium was analyzed offline daily by Protein A chromatography on an Agilent Bioinert 1260 HPLC system using a Bio-Monolith Recombinant Protein A column (Agilent Technologies, Santa Clara, CA) as well as with the BLItz biolayer interferometer system (ForteBio, Fremont, CA) using a Protein A biosensor (ForteBio, Fremont, CA).

### Perfusion Bioreactor Culture

The perfusion cell culture experiments were conducted in a 5 L glass bioreactor using a Biostat B-DCU controller (Sartorius, Göttingen, Germany) with a working volume of 2 L. Temperature and pH control was initiated before inoculation to 37 °C and 7.1, respectively. Dissolved oxygen was controlled to a setpoint of 50%. The pH was controlled by either sparging CO_2_ or bolus additions of 0.5 M NaOH (Sigma Aldrich, St. Louis, MO). The bioreactor was inoculated to an initial density of 0.5 × 10^6^ cells/mL. An XCell™ ATF 2 stainless steel device (Repligen, Waltham, MA) with a 0.2 μm hollow filter fiber cartridge (Repligen, Waltham, MA) controlled by an XCell C24 Controller (Repligen, Waltham, MA) was used for steady-state perfusion, slowly ramping up the exchange rate from 0.25 to 1.0 vessel volumes a day (VVD) between days 4 and 8. The culture was then maintained at 1 VVD thereafter until harvest day. The bleed rate was adjusted proportionally with the permeate rate to maintain a constant VVD and viable cell density (VCD) throughout the culture duration. Offline samples were taken daily to analyze various culture parameters as described above. Titer and glycans were analyzed at-line daily using the N-GLYcanyzer PAT system as previously described.^28^

### N-Glycosylation Analysis

Offline N-glycan analysis was done using AdvanceBio Gly-X N-glycan prep with InstantPC (GX96-IPC, Agilent Technologies, Santa Clara, CA) following the manufacturer’s instructions. Briefly, spent media was removed from the bioreactor daily and then purified using a Protein A HP SpinTrap (Cytiva, Marlborough, MA) with 20 mM phosphate buffer pH 7.2 as a binding buffer and 0.1% formic acid as the eluent. The sample was then neutralized using 1 M HEPES solution pH 8.0 before buffer exchange into 50 mM HEPES solution pH 7.9 and then concentrated to ∼ 2 g/L using a 10 kDa MWCO spin column (VWR, Radnor, PA). After, 2 mL of Gly-X denaturant was added to 20 mL of the sample before heating to 90 °C for 3 min. After cooling, 2 mL of N-Glycanase working solution (1:1 Gly-X N-Glycanase:Gly-X Digest Buffer) was added, mixed, and incubated at 50 °C for 5 min. Afterward, 5 mL of Instant PC Dye solution was added, mixed, and incubated for an additional 1 min at 50 °C. The sample was diluted with 150 mL of load/wash solution (2.5% formic acid, 97.5% acetonitrile (ACN)). Then, 400 mL of load/wash solution was added to the Gly-X Cleanup Plate along with the ∼ 172 mL of sample. A vacuum was applied (<5 inHg) until the sample passed through. Samples were washed twice with 600 mL of Load/Wash solution before being eluted into a collection plate with 100 mL of Gly-X InstantPC Eluent with vacuum (<2 in Hg). The at-line sample preparation for perfusion follows the same scheme and is described in detail in an earlier publication.^29^

These samples were run on a 1260 Infinity II Bio-Inert LC System (Agilent Technologies, Santa Clara, CA) using an AdvanceBio Glycan Mapping column 2.1 × 150 mm 2.7 micron (Agilent Technologies, Santa Clara, CA). Mobile phase A was 50 mM ammonium formate adjusted to pH 4.4 using formic acid and mobile phase B was ACN. The flow rate was set to 0.5 mL/min, the FLD was set to ex. 285 nm/ em. 345 nm, and the column temperature was set to 55 °C. The initial eluent was held at 80% B for 2 mins, then dropped immediately to 75% B. From 2 to 30 mins the eluent was changed from 75% B down to 67% B in a linear gradient, and then from 30 to 31 mins, it was decreased from 67% B down to 40% B. From 31 to 33.5 minutes, the ACN concentration was brought back to 80% at which level it was held until the end of the run at 45 mins.

Relative estimation of glycan fractions was done on OpenLab CDS v3.5 (Agilent Technologies, Santa Clara, CA). Relative accumulated abundances were calculated by integrating each glycan peak under its respective curve. Each glycoform was then calculated by dividing the respective glycan peak area by the total peak area. Accumulated galactosylation Index (GI) was calculated by the summation of all galactoforms from all abundant glycoforms. Accumulated Mannosylation Index (MI) was measured similarity to galactosylation index but by summating the abundance of all mannosylated glycoforms. The daily temporal changes in each glycoform were calculated by calculating the change in mAb titer and relative abundances required between time points to reach the glycan fraction for the next time point (day).

### Amino Acid Preparation and Analysis

Extracellular amino acids were measured using an Agilent AdvanceBio Amino Acid reagent kit and a 3 mm I.D. AdvanceBio AAA C18 column following the manufacturer’s instructions. Briefly, 200 μL of supernatant from each daily bioreactor sample was taken and placed onto the HPLC multi-sampler in a vial. The multisampler needle individually prepared samples prior to injection. First, 2.5 μL of borate buffer was aspirated from a vial, followed by 1 μL of the sample, and the two were mixed before waiting 12 s. Afterward 0.5 μL of ortho-phthalaldehyde (OPA) was aspirated from a vial and mixed before aspirating 0.4 μL of Fluorenylmethyloxycarbonyl chloride (FMOC) from a vial and the two were mixed. Then the sample was diluted by aspirating and mixing 32 μL of injection diluent (0.4% concentrated H_3_PO_4_ in mobile phase A). The complete sample is then injected for analysis. Detection was done using a DAD system using two signals: Signal A at 338 nm, reference wavelength 390 nm for OPA-derived amino acids and Signal B at 262 nm with reference wavelength at 324 nm for FMOC-derived amino acids. Mobile Phase A was 10 mM Na_2_HPO_4_, 10 mM Na_2_B_4_O_7_ pH 8.2, and mobile phase B was 45:45:10, v:v:v ACN:methanol:water. The column was set to 40 °C and a flow rate of 0.6 mL/min.

### Nucleotides Sugar Extraction and Analysis

The intracellular nucleotide sugars were extracted from CHO Cells following a method by Sha et al.^30^ Briefly, a volume of 3 million cells was collected and centrifuged at 1000 RPM for 5 minutes. The supernatant was discarded. The cells were resuspended with 1 mL of cold PBS as a wash before being centrifuged again. The PBS was discarded, and the pellet was flash-frozen on dry ice and stored at -80 °C until extraction. For extraction, cell pellets were thawed and then 200 μL of 0.5 M PCA was added to disrupt the cells and allow nucleotide sugars to release. The sample was spiked with 0.5 μL of 20 mM GDP-Glc as an internal standard. The lysate was incubated on ice for 5 mins before being centrifuged at 2,000 xg for 3 mins at 4 °C. The supernatant was removed and transferred to a new tube. The cell pellet was then again subjected to 200 μL of 0.5 M PCA and 0.5 μL of 20 mM GDP-Glc and incubated on ice for 5 mins before being centrifuged at 18,000 xg for 3 mins at 4 °C. The two supernatants were then merged and neutralized with 56 μL of 2.5 M KOH in 1.1 M K_2_HPO_4_. The sample was then incubated for 2 more mins on ice before being centrifuged at 18,000 xg for 1 min to remove the potassium perchlorate precipitate. The supernatant was filtered with a 0.2 μm PVDF syringe filter into a new tube and stored at 4 °C until HPLC analysis. A 5-point calibration curve was made for all nucleotide sugars for quantitation.

### Data Analyses

The change in mAb production was calculated by equation 1 (eq. 1), by dividing the change in titer between two time points, mAb titer at time 2 (t2) and mAb titer at time 1 (t1). The integrated viable cell density (IVCD) was calculated by equation 2 (eq. 2), where the change in viable cell density (VCD) was calculated by the summation of the VCD at two time points multiplied by the change in time, divided by 2. Productivity was measured by equation 3 (eq. 3), where the productivity is equal to the change in mAb titer divided by the change in IVCD between two time points. This equation was also used to measure consumption rates of glucose and galactose by swapping the change in mAb titer for either change in glucose or galactose concentration.

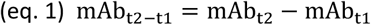

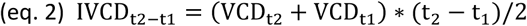

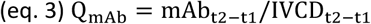

## Results & Discussion

### Fed-Batch Culture Performance

This work aims to understand how the addition of galactose as a supplemental carbon source may influence mAb production and N-linked glycosylation over time. Here we looked at the addition of a feed, either glucose or glucose with galactose, while also varying the feeding times to be either daily (24 h) or alternative days (48 h) to understand how this may influence the CHO cell culture as well as the mAb that is produced. Manganese was added to the galactose-supplemented cultures to 1 μM during the first feeding to increase relative glycosylation further.

The viable cell densities (VCD) between all conditions influenced the peak viabilities and the longevity of all cultures that were run. Cultures fed daily had a higher peak density than those fed only on alternative days (**Figure 1A)**. Daily feeding control cultures using both feed strategies saw a peak VCD of 26.6 × 10^6^ cells/mL, while alternative day control and supplementation cultures saw a peak VCD of 22.8 × 10^6^ cells/mL and 24.3 ×10^6^ cells/mL, respectively. A paired T-test showed statistical significance between the feeding times (daily vs. alternative day) but not between the control and supplementation groups (**Supplemental Figure S1**). Based on these results, the timing of feeding can be inferred as having a more considerable impact on the peak VCD than the addition of galactose as a secondary carbon source. When looking at culture longevity (**Figure 1B)**, all cell culture conditions survived with cell viability above 70% up until day 10 and were harvested once the cell viability fell below this threshold. Only one condition, the alternative day-fed culture with supplementation, survived until day 11. Because of this, we decided to use day 10 as the cutoff for all further analyses within this study.

**Figure 1:**
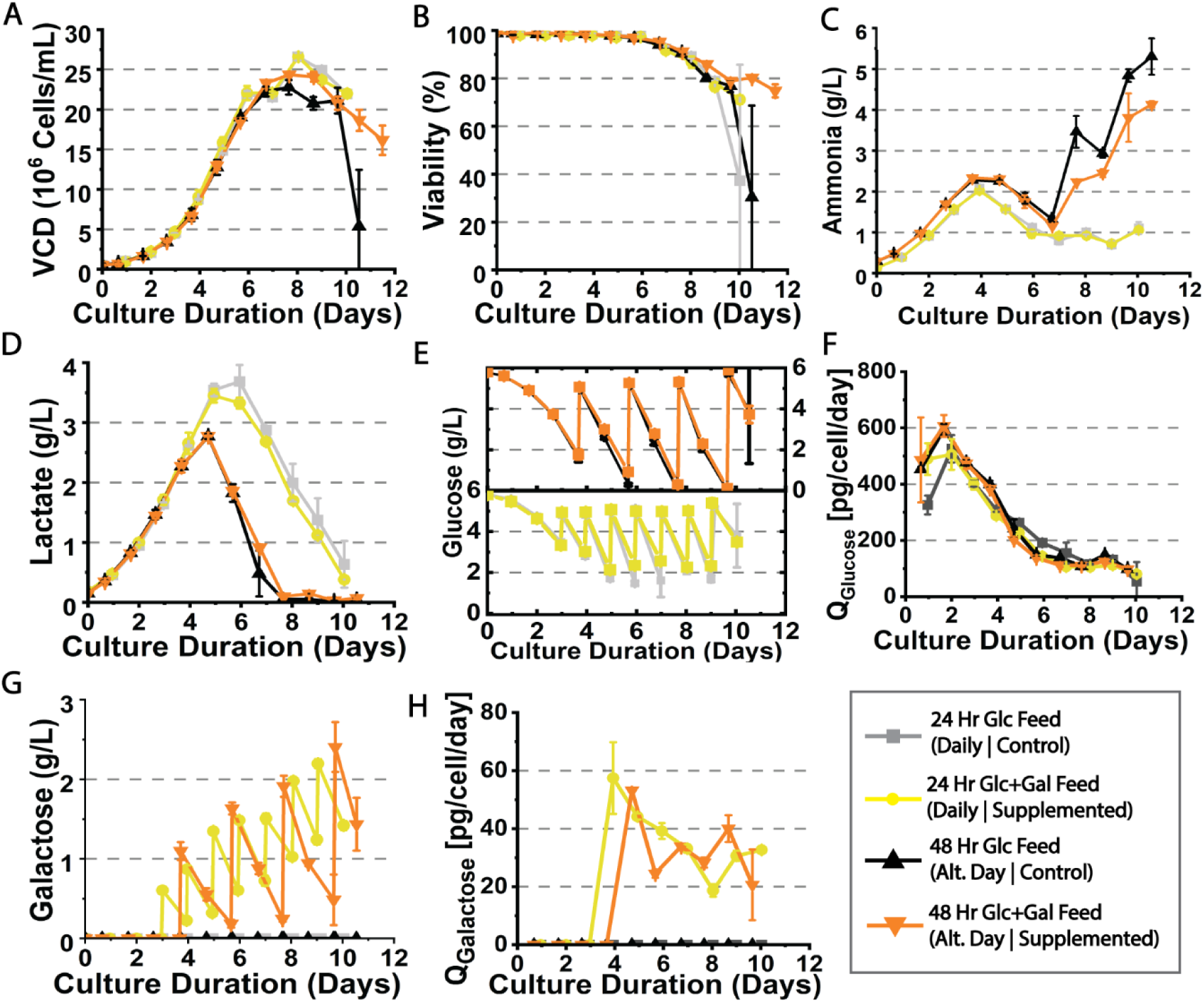
Summary of fed batch culture process performance metrics. Viable cell density (A), viability (B), ammonia content (C), lactate content (D), glucose content (E), glucose consumption rate (F), galactose content (G) and galactose consumption rate (H).

Ammonia production followed similar trends between feeding regimes until day 7, when trends diverged based on the feed frequency (**Figure 1C**). The cultures fed daily saw a peak ammonia concentration around day 4 whereas the control and supplemented cultures both peaked at ∼2 g/L ammonia. Afterward, the ammonia concentration in these cultures decreased to roughly 1 g/L. No statistical significance was seen between these groups. For the alternative day control and supplementation cultures, ammonia peaked on day 4 at 2.23 and 2.31 g/L, respectively. The ammonia concentration then decreased until day 7 and rapidly increased until day 10 where the alternative day feed control saw a peak on day 10 of 5.31 g/L ammonia, while the alternative day supplementation saw a peak of 4.13 g/L. The increase in ammonia concentrations signals metabolic stress within the culture and indicates shifts in amino acid metabolism, especially within alanine metabolism.^31^ Extracellular alanine concentration peaked at 3 mM for the alternative day feeding cultures and then decreased until the end of the culture. However, for daily fed cultures, the extracellular alanine concentration increased throughout the culture until peaking on day 10 at 5.5 and 4.1 mM alanine for daily control and supplemented cultures, respectively (**Supplemental Figure S2**). One explanation could be a more efficient utilization of ammonia detoxication of the daily fed cultures through alanine production within these cultures by the conversation of intracellular ammonia and pyruvate into alanine.^31^

Lactate production (**Figure 1D**) followed a similar trend between feeding days, reaching a peak lactate concentration on day 5 before the metabolic shift to lactate consumption. The daily control and supplemented feeds both saw a peak lactate production of 3.5 g/L. Lactate concentration in the medium then decreased through the rest of the culture. Alternative day feeding cultures saw a lower peak lactate concentration on day 5 of 2.77 g/L. These cultures then saw a decrease in the lactate levels until day 8 and stayed low until day 10. The increased lactate production can be attributed to the more periodic feeding rate of the glucose allowing for higher glucose metabolism towards lactate.^32^ For this reason, the alternative day-feeding cultures saw lower glucose levels than the daily-fed counterpart, signaling a metabolic switch within the cells.^33^

## Fed-Batch Consumption of Glucose and Galactose

To explore the influence of glucose and galactose consumption, the cultures were controlled to 5 g/L glucose daily or on alternative days, as explained in the methods section. To characterize galactose as a feed additive, the feeds were spiked with galactose at a ratio of 3:10 (galactose: glucose w/w). This ratio was decided based on past literature examining galactose dose dependence’s influence.^17,34,35^ Similarly, the galactose-supplemented cultures were spiked to 1 μM manganese on the first feed day, a dose chosen to mirror previous literature.

The top graph of **Figure 1E** shows the extracellular glucose present for the alternative day feeding regimens, while the bottom graph shows glucose concentrations for daily feedings. For the alternative day feeding regimen, the first feeding was done on day 4 when the glucose concentration was 1.63 g/L for the control and 1.77 g/L for the supplemented cultures. The glucose within the cultures was depleted on feeding days after the first feed. Before the second feed, the glucose concentrations were 0.33 g/L (control) and 0.92 g/L (supplemented); the differences between these concentrations may be attributed to having galactose as a secondary carbon source within the supplemented cultures. There was no difference in glucose consumption trends for the alternative day feeding regimens based on glucose consumption rates **(Figure 1F)**. However, it may be noticed that the alternative day feeding regimens did not vary largely from one another, and consumption rates tended to be higher during the first 4 days of the culture versus the daily feeding cultures. For the daily feeding samples, the first feed was done on day 3 when the concentrations were around 3.5 g/L for the control and 3.3 g/L for the supplemented cultures. The difference in glucose concentrations within these cultures’ conditions can be seen and shows that there was less consumption of glucose when galactose was present in the culture. However this may be caused by the co-consumption of lactate during the latter phase of the cultures on and after day 5. Nevertheless, results are in agreement with earlier work looking at the comparison of glucose consumption in the presence of galactose.^21^

There was a steady accumulation of galactose in the supplemented cultures (**Figure 1G**). Specific galactose consumption rates were highest within the first 24 hours of supplementation and would decrease over time **(Figure 1H)**. This may be caused by the drastic increase in the concentration after the initial feeding of galactose. Both supplemented culture conditions saw a consumption rate between 20 and 40 pg/cell/day. Interestingly, the daily supplemented cultures saw a slow plateau in the consumption rates of galactose while the alternative day cultures had an increased rate after feeding and lower rates on days in between feeding. These changes in the specific consumption rate are consistent with previous studies where the galactose consumption rate is dose-dependent and higher galactose in the medium will increase the consumption rate until a point of saturation.^21,36^

## Fed-Batch Culture Productivity

MAb titer and productivity were studied to understand the influence of the feeding regimen and supplementation. All cultures reached their peak titer around day 9, then saw a titer decrease within the reactor on subsequent days (**Figure 2A**). This loss may be due to protein degradation. Daily feeding cultures saw a peak titer of 1.12 g/L (control), and 1.23 g/L (supplemented). The alternative day feeding samples saw a peak titer of 1.11 g/L (control) and 1.19 g/L (supplemented). A paired comparison plot analysis did not show any statistical significance between these cultures in respect to peak titer (data not shown). This may be due to a nutrient limitation as extracellular asparagine was exhausted by day 6 in all cultures **(Supplemental Figure S2)**, a critical amino acid known to influence mAb production in CHO Cells.^37–39^

**Figure 2:**
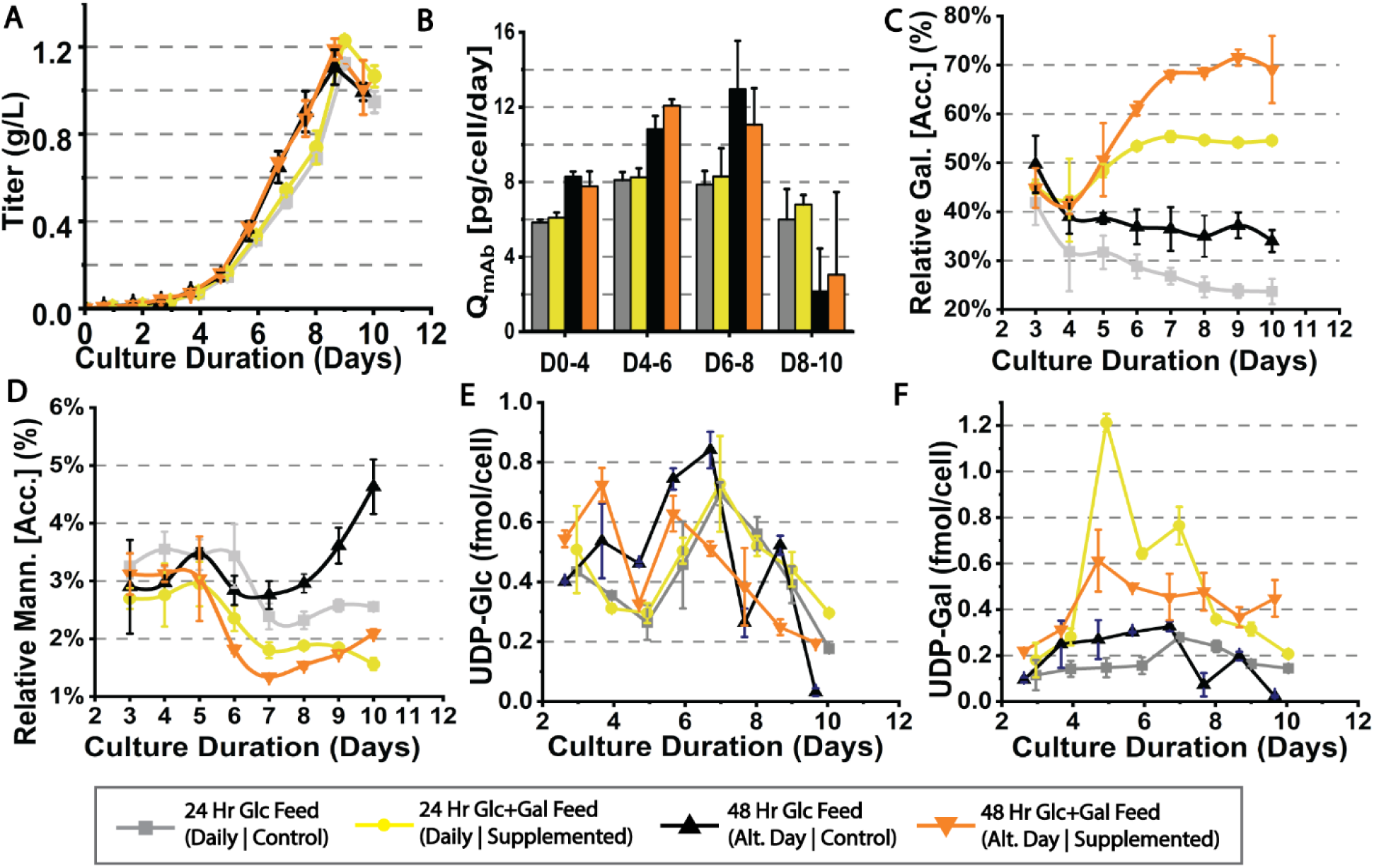
Productivity, glycan indices, and glycan precursors for fed batch cultures. MAb titer (A), mAb specific productivity (B), relative accumulated galactosylation (C), relative accumulated mannosylation (D). The intracellular nucleotide sugar pools of UDP-Glucose (E) and UDP-Galactose (F) are presented here, which are precursors for mAb glycosylation.

When comparing cell-specific production rates (**Figure 2B**), a trend can be seen between the feeding regimens. The average cell-specific mAb productivity during the first 6 days of culture was higher for the alternate-day feeding cultures (control and supplemented) versus that of the daily feeding cultures. However, there was no statistical difference between the control and supplementation groups, meaning that the addition of galactose and manganese did not influence the mAb productivity rate, and any minor difference in productivity is attributed to the feed timing. Since the alternative day cultures were controlled to 5 g/L glucose, glucose depletion did not interrupt mAb productivity. Fan *et al*. investigated the influence of mAb productivity under glucose starvation and found an inverse correlation between mAb productivity and the degree of glucose starvation. However, mild starvation did not influence mAb productivity, which is in alignment with the results seen in our study.^40^

## Fed-Batch Monoclonal Antibody Glycan Indexes

Two important glycan indices, galactosylation index (GI, **Figure 2C**) and mannosylation index (MI, **Figure 2E**) were calculated from the relative abundances of all observed glycoforms. These values are accumulated relative abundances per day. A separate graph for each glycoform is shown in **Supplemental Figure S3**.

A truncated explanation of the interconnected network and metabolism between glycolysis, the Leloir pathway and glycosylation is portrayed in **Scheme 2**. Glucose is taken up by the cell and used for glycolysis, where part of the energy produced goes toward the TCA cycle and protein production. Galactose is also taken up by the cell and metabolized by the Leloir pathway, where it can then enter the glycolytic pathway via the intermediate glucose-1-phosphate (Glc-1-P). However, part of the incoming galactose will also produce UDP-galactose (UDP-gal), a glycan precursor to produce galactosylated mAb species. The bottom of the scheme briefly depicts how the glycosylated mAb is shuttled from the endoplasmic reticulum (ER) to the Golgi apparatus while going through multiple trimming steps from Man8 to eventually G0 or G0F after the addition of GlcNac to the chitobiose core. From there these transient species can become galactosylated by the addition of galactose by one of several galactosyltransferases (i.e., B4GALT) using manganese as the cofactor.

MAb galactosylation on day three did not show any statistical significance between the cultures as the sampling was done prior to the first feed addition. Interestingly, the GI followed similar trends between the galactose-supplemented and control cultures, where the daily feed control had a GI of 23.7% on day 10, roughly 10% less than the alternative day feeding control, which had a GI of 34%. Cultures fed with galactose and spiked with manganese saw a significant increase in the rate of galactosylation against their glucose-fed control counterparts. Daily fed cultures supplemented saw a GI of 55.5%, while the alternative day-supplemented feeding samples had a GI rate of 69.1%, a 13.6% increase. The alternative day feeding strategies increased the rate of galactosylation more than their daily feed counterparts. Galactose-supplemented cultures saw an increase in GI as this allowed for an increase in UDP-Gal biosynthesis from the fed galactose through the Leloir pathway.^41^ There is a positive correlation between UDP-Gal levels (**Figure 2F**) and the relative GI within the first half of the cultures’ duration. Although the mAb productivity was higher with the alternative day feeding regimen (**Figure 2B**), the increased UDP-Gal pool and a possible increase in the galactosyltransferase activities may aid galactose occupancy of the produced mAbs. These findings agree with Fan et al.’s similar glucose starvation study, where high and severe glucose starvation cultures saw an increase in GI versus low starvation and non-glucose starved cultures.^40^

Relative accumulated mannosylation is shown to be under 3% for all cultures besides the alternative day feed control (**Figure 2D**). With the addition of galactose and manganese to the cultures, the MI index decreases for both feeding regimens within the study to 2% for the alternative day supplementation cultures and 1.6% for the daily feed-supplemented cultures. Without the addition of galactose and manganese, the daily feed control culture saw a MI of 2.5% while the alternative day feeding control saw a statistically significant MI of 4.5%. This increase in mannosylation is due to the high ammonia accumulation in the alternative day control culture and is usually associated with low UDP-GlcNac pools.

The UDP-glucose (UDP-glc) pool (**Figure 2E**) within the alternative day feeding control saw a spike in levels after each feeding. However, UDP-glc levels quickly decreased after day 7, along with the UDP-gal levels. This may indicate that some of the nucleotide sugars are metabolized towards other pathways, instead of towards glycosylation. This is also further exemplified when comparing the nucleotide sugar pools and MI for the alternative day feed control versus the daily feed control cultures. Low glucose concentrations in the medium are known to deplete precursors required for glycan maturation which can also be linked with the UDP-glc levels.^42–44^

## Fed-Batch Temporal Changes in Glycosylation

To build a temporal understanding of the glycosylation within the fed-batch cultures, the daily relative glycan isomers were calculated based on daily accumulation rates and titers based on changes in weighted averages (**Figure 3)**. The values were calculated until peak titer was reached (culture day 9) since subsequent days saw a drop in titer. Sialylated species are not shown as there were minimal to no changes in these species regarding daily glycosylation rates. Each glycoform showed a similar trend as with the accumulated glycan index values mentioned above, where cultures supplemented with galactose and manganese saw an increase in galactosylated species and a decrease in other transient and truncates species such as G0F-GlcNac and G0F.

**Figure 3:**
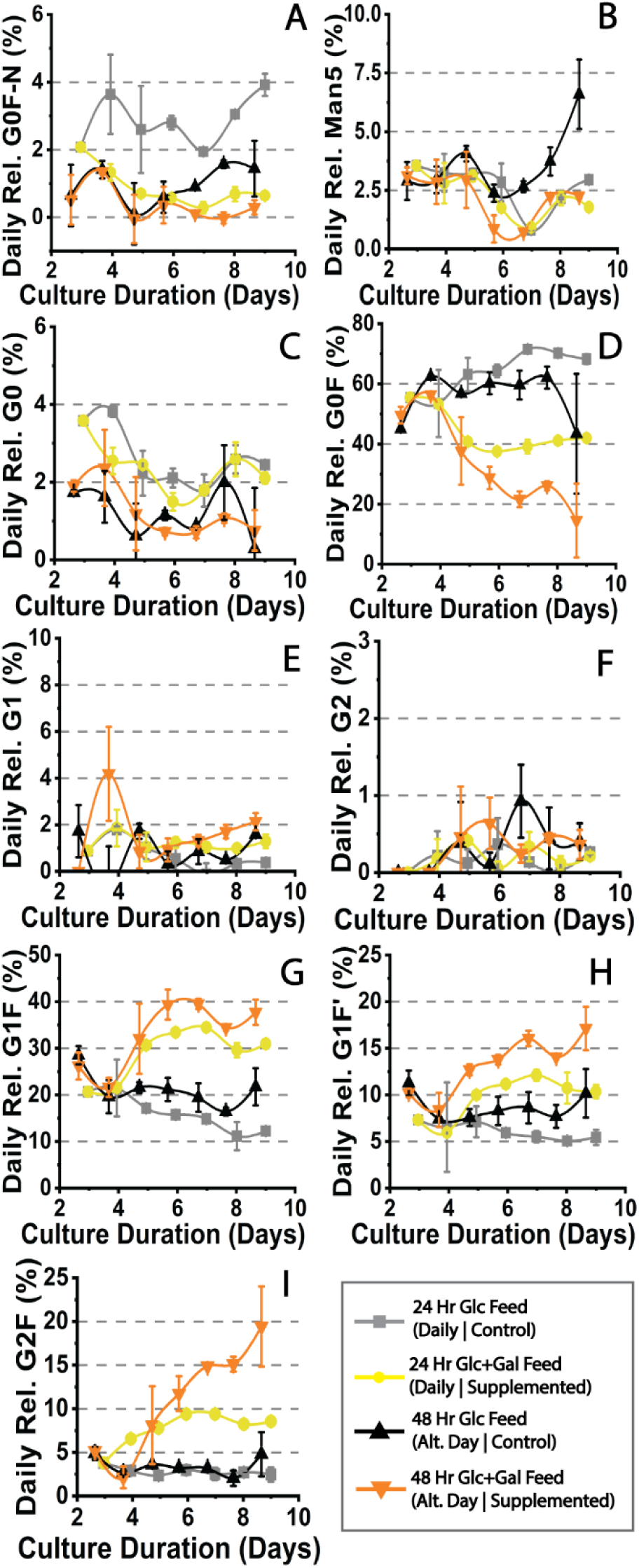
Relative daily glycoform production for fed-batch cultures. The major mAb glycoforms are displayed here. Sialylated glycoform species are omitted as there was little to no change in their output. Their data are provided in Supplemental Figure S3.

When looking at the truncated G0F-GlnNac isoform (**Figure 3A**), this species tended to stay between 1-2% under all conditions besides the daily feed control culture, where the value increased to roughly 4%. Similarly, for the G0 isoform (**Figure 3C**), the abundance tended to decrease through culture duration. However, the alternative day feeding regimen tended to decrease the abundance of this isoform more than daily feed cultures.

G0F tends to be the most abundant glycoform seen for mAbs (**Figure 3D**). There is an increase in this glycoform under the control daily feeding regimen compared to that of the alternative day control, where the relative daily abundance of this isoform is lower through the second half of the cultures. A bottleneck in processing this glycoform further to produce galactosylation is assumed as the G0F glycoform tends to increase over time in the culture. When galactose and manganese are added, there is a decrease in this bottleneck. With daily supplementation culture, the G0F glycoform reaches a steady state secretion of G0F glycoforms around 40% at day 5. However, when compared with the alternative day supplementation cultures, a steady decrease in the G0F glycoform is seen, and on day 9 the relative abundance of the G0F glycoform produced is under 20%. This would indicate that under the alternative day feeding, where glucose is starved in the culture, the metabolism of galactose is toward producing galactosylated species of mAb than toward energy metabolism.

When looking at the galactose mAb isoforms, new information regarding the galactosylation rates can be revealed. This is not very apparent when looking at the lower abundance galactoforms such as G1 (**Figure 3E)**, G1’ (not shown), or G2 (**Figure 3F**), as small changes are harder to capture with these isoforms. **Figure 3G and 3H** show G1F and G1F’ glycoforms. An inverse relationship can be seen here versus the G0F isoform as a decrease in the G0F isoform and an increase in the G1F+G1F’ isoforms indicates that the addition of galactose and manganese increased the galactosylation of the antibody. The cultures without supplementation saw a significantly lower, but steady production rate of the double galactose occupied glycan isoform G2F (**Figure 3I**). During daily supplementation, there was an increase of the G2F isoform that produced a relatively stable abundance of G2F throughout the second half of the culture, around 8-10% relative G2F species a day. In comparison, the G2F species increased in its daily produced abundance and reached almost 20% by day 9 when the culture was subjected to the alternative day supplementation feeding regimen. This can help understand why a discrepancy is seen in the GI between the alternative and daily supplementation samples, as the alternative day-supplemented cultures had an increase in the production of the G2F glycoform over time. However, since sialylation was not altered with the supplementation of feeding interval, there may be another bottleneck with the corresponding sialyltransferase activities. Though, manganese and galactose supplementation does not influence the activity of sialyltransferase and Trastuzumab is known to have little to no sialic acid glycoforms.

It is also worth mentioning that the relative daily abundances of the high mannose isoform Man5 (**Figure 3B**) was not influenced by the addition of galactose or manganese. When comparing the daily and alternative day feeding regimen controls, the daily fed cultures saw a Man5 abundance similar to the supplemented cultures (between ∼2-3%). However, the alternative day feeding control saw an increase in Man5 on and after day 7 of culture and peaked by day 9 at ∼6.5%. This increase in the high mannose species indicates stress within the glycosylation metabolism. One hypothesis is that the glucose metabolism was toward energy production rather than mAb glycosylation as the extracellular glucose levels were critically low by day 5 of the cultures with alternative day feeding regimens.

## Perfusion Culture Performance

Perfusion cultures were run to further elucidate the glycosylation dynamics where there were no limitations to nutrients as in fed-batch. These cultures were run similarly, where the control used HIP-CHO medium, and the supplemented medium consisted of HIP-CHO medium (containing 6 g/L glucose) with the addition of 1.8 g/L galactose and 1 μM manganese. This glucose to galactose ratio is equivalent to the feed ratio used in the fed-batch study. Both cultures were run as batch cultures until the glucose concentration in the reactors fell to or below 2 g/L, at which time perfusion was started to increase nutrients in the bioreactor culture. The permeate and bleed pumps were manually adjusted daily to reach and maintain a cell density of around 20 million cells/mL under both conditions (**Figure 4A**). The pumps were adjusted proportionally to maintain 1 VVD throughout steady-state perfusion. Experiments were compared to day 13; however, the control reactor was run for 20 days.

**Figure 4:**
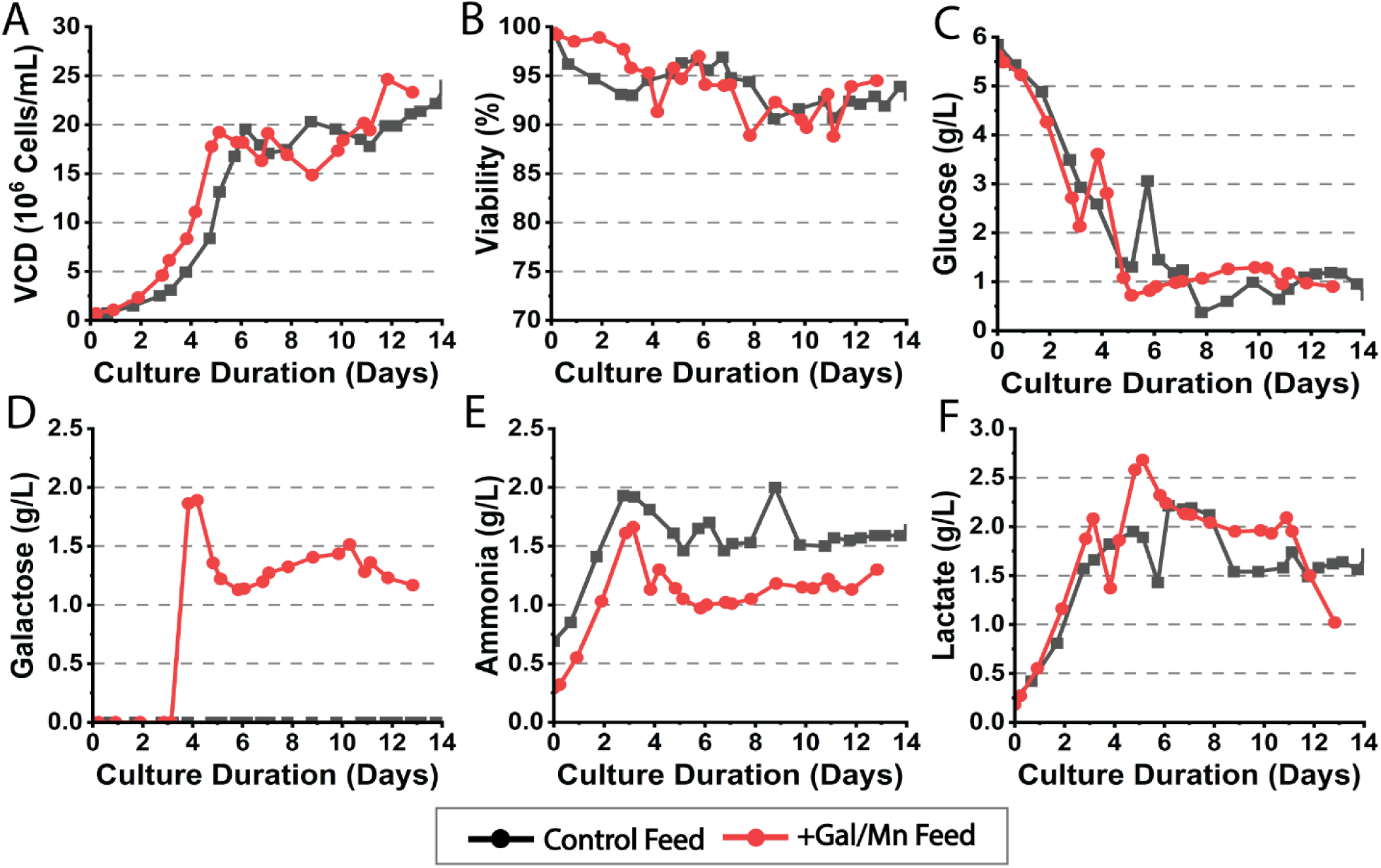
Perfusion culture process conditions and metrics. Viable cell density (A), viability (B), glucose content (C) galactose content (D), ammonia content (E) and lactate content (F).

For the process conditions, the cultures’ viability (Figure 4B) under both conditions stayed above 90% for most of the run. Glucose levels in the culture were maintained roughly between 0.5 and 1 g/L under both conditions (**Figure 4C**). The galactose levels (**Figure 4D**) in the +Gal/Mn supplemented culture saw a galactose level within the medium between 1 and 1.5 g/L, which would indicate around 0.3 – 0.8 g/L of galactose was being consumed daily during steady-state perfusion after reaching peak cell density. Ammonia levels within the +Gal/Mn supplemented culture were lower than that of the control by roughly 0.5 g/L (**Figure 4E**), and lactate levels were higher on average for the +Gal/Mn supplemented culture versus the control (**Figure 4F**).

When comparing the extracellular amino acid profiles between culture conditions (**Supplemental Figure S4**), the amino acids all followed a similar trend, however, the +Gal/Mn culture had a higher level of aspartate in the culture which is correlated with higher cell productivity but not with cell growth.^45^ These results are consistent with the increased mAb productivity, as explained in the next section.

## Perfusion Culture Titer and Glycosylation

On average, the +Gal/Mn supplemented bioreactor produced more product versus the control run (**Figure 5A**). Perturbations can be seen in each condition which can be accounted for by manual changes in the bleed and permeate pumps which influence the cell density. However, even with these perturbations, the reactor titer for the +Gal/Mn supplemented reactor produced more product than the control. Under fed-batch experimentation, it is known that the use of galactose as a substitute for glucose will have diminishing returns on production due to the lower transport of galactose over glucose. A study by Gramer et al. looked at the use of uridine, manganese and galactose on a fed-batch culture and saw a low bolus addition of these feed additives (i.e., 0.9 g/L galactose, 2 μM MnCl_2_, and 1mM uridine) will have a positive impact on titer, but will marginally decrease with an increase in galactose, manganese and uridine concentrations.^21^ This may also apply to perfusion, which may give insight into why the productivity is higher than the control culture. To our knowledge, no literature relating the addition of galactose as a feed supplement to productivity increase exists. However, literature that looks to substitute glucose partially with galactose is known to have an unfavorable influence on productivity and final titer.^46,47^

**Figure 5:**
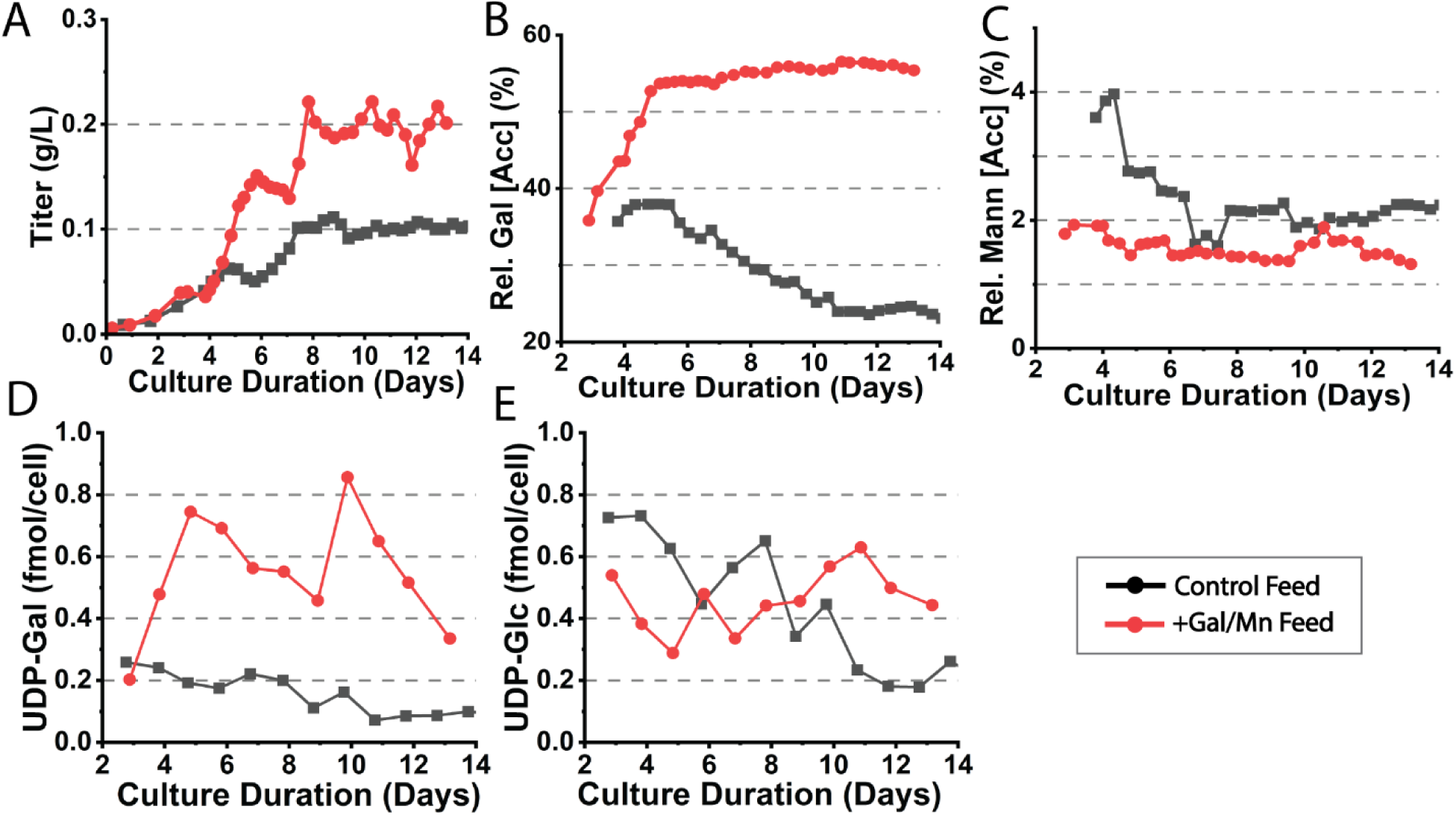
Titer, glycan indices, and glycan precursors for perfusion cultures. Daily reactor titer (A), relative galactosylation index (B) and mannosylation index (C) are shown. The cell-specific concentration of nucleotide sugar glycan precursors UDP-galactose (D) and UDP-glucose (E) are shown as well.

The GI can be seen in **Figure 5B**. The galactosylation rate for the control culture was similar to the 24-hour control fed-batch culture in which the relative galactosylation reached around 25% by day 10. For the +Gal/Mn supplemented culture, the galactosylation rate increases to 54-56% through the culture duration, similar to the 24-hour supplement-fed batch culture experiment described above. Investigating the UDP-Glc levels (**Figure 5E**), fed-batch cultures saw a decrease of the UDP-Glc levels toward the end of the culture for the control cultures. A similar trend can be seen with perfusion; however, the control perfusion run never depletes its UDP-Glc pool since there is a constant influx of glucose into the medium. The +Gal/Mn perfusion culture exhibited a constant UDP-Glc level within the cells, interestingly after day 10 the control saw a drop in the UDP-Glc levels while the level stayed between 0.4-0.6 fmol/cell with the +Gal/Mn culture. This could be accounted for by the conversion of UDP-Gal to UDP-Glc.

When comparing the UDP-Gal levels between the perfusion cultures, a significant increase can be seen by adding galactose and manganese to the feed (**Figure 5D**). This high level of UDP-Gal in the +Gal/Mn explains the increasing rate of galactosylation. Here it seems that the highest level of galactosylation is 55%, while under fed-batch alternative day feeding where glucose was starved a higher accumulated rate of 70% was seen. This is an interesting phenomenon, as described earlier. Running a perfusion reactor with continuous feeding rather than daily or alternative day feeding seems to have a lower rate of galactosylation. Further studies can be done to understand the influence of glucose starvation on galactosylation to elucidate further pathway changes that influenced a favorable increase in the galactosylation rate.

The MI (**Figure 5C**) also saw a steady state of mannosylation at around 2% for the control culture and 1.5% for the +Gal/Mn culture. These results are similar to our results with the fed-batch daily feeding control and Gal/Mn-supplemented cultures. A breakdown of each glycoform present can be seen in **Supplementary Figure S3**.

## Conclusion

In this work we have sought an understanding of the temporal changes of mAb glycosylation patterns during CHO cell culture bioprocessing explained by changes in the key metabolites when feeding galactose and manganese into the cultures. This was accomplished by studying two operation modes: (1) a fed-batch bioprocess in which galactose was fed proportionally with the glucose feed, as well as with the addition of manganese on the first day of feeding, either daily or on alternative days to understand glucose limitation as a secondary variable in the study; (2) a perfusion operation (non-nutrient limited), where galactose and manganese were spiked into the media and the culture was maintained in a steady state.

For the fed-batch study, results showed that the feeding schedule, rather than the addition of galactose, had a profound and favorable effect on peak cell density. Daily feeding regimens increased the peak lactate produced by 0.7 g/L over alternative feeding strategies. However, alternative day feeding saw higher ammonia production in the latter stages of the culture as daily fed cultures were better able to metabolize ammonia to alanine. Galactose feeding and feeding times had a marginal impact on the glucose consumption rate, which may also be due to co-consumption of lactate. Galactose consumption rates were found to increase with the extracellular galactose concentration, consistent with previous findings. While galactose and the feeding regimen did not influence peak titers, the alternative day fed cultures saw higher productivity on average through the first half of culture. In addition to these differences in culture performance, the feeding regimens were found to have the most considerable impact on mAb glycosylation. The addition of galactose and manganese significantly increased the rate of galactosylation with alternative day fed cultures exhibiting the highest relative galactosylation of 70% compared to 50% of the daily galactose fed cultures. Mannosylation also decreased in all instances with the galactose and manganese additions over the controls.

For the perfusion study, the addition of galactose and manganese increased the titer over the control culture. An increase in the galactosylation was also observed where the galactose and manganese-supplemented culture saw 55% relative galactosylation, while the control decreased to roughly 25%. The addition of galactose was able to restore UDP-glucose pools over the control where UDP-glucose was depleted over the duration of the culture. Overall, the perfusion study gave more insight into the glycosylation network and the influence that glycan precursors have on metabolism and the glycosylation pathway.

In summary, we were able to gain a temporal understanding of how galactose and manganese influence mAb product quality. Future work will focus on developing the understanding and PAT necessary to build process control schemes based on more precise dosing of glycan precursors to promote homogenous glycosylation during upstream bioprocessing.

## Disclaimers

The authors declare that they have no competing financial interests. This article reflects the author’s views and should not be construed to represent FDA’s views or policies. Certain commercial equipment, instruments, or materials are identified in this paper to foster understanding. Such identification does not imply recommendation or endorsement by the FDA.

## Author Contributions

### Aron Gyorgypal

Conceptualization, Investigation, Methodology, Data Curation, Formal Analysis, Writing – Original draft, Writing: review & editing. **Erica Fratz-Berilla:** Conceptualization, Investigation, Methodology, Data Curation, Supervision, Writing: review & editing. **Casey Kohnhorst:** Conceptualization, Investigation, Methodology, Data Curation, Supervision, Writing: review & editing. **David N. Powers:** Conceptualization, Investigation, Methodology, Supervision, Writing: review & editing. **Shishir P.S. Chundawat:** Conceptualization, Writing - Review & Editing

## Acknowledgments

This work was supported by NIIMBL Project Award PC5.2-112 and supported in part by the appointment of Aron Gyorgypal to the Research Participation Program at FDA, administered by ORAU through the US Department of Energy Oak Ridge Institute for Science and Education (ORISE). The authors would like to thank the Office of Biotechnology Products (OBP) Bioprocessing Lab (Dr. Cyrus Agarabi, Dr. David N. Powers, Ms. Xin Bush, and Ms. Nicole Azer) at the US FDA Center for Drug Evaluation and Research (CDER) for their support in this project. The authors also thank Dr. Natasha Ilyushina (CDER/OBP) and Dr. Omnia Ismaiel (CDER/Office of Clinical Pharmacology (OCP)) for their critical review of this manuscript. The authors also thank GenScript Biotech Corporation (Piscataway, NJ) for the Trastuzumab cell line gift to Rutgers University.

## Scheme

**Scheme 1:**
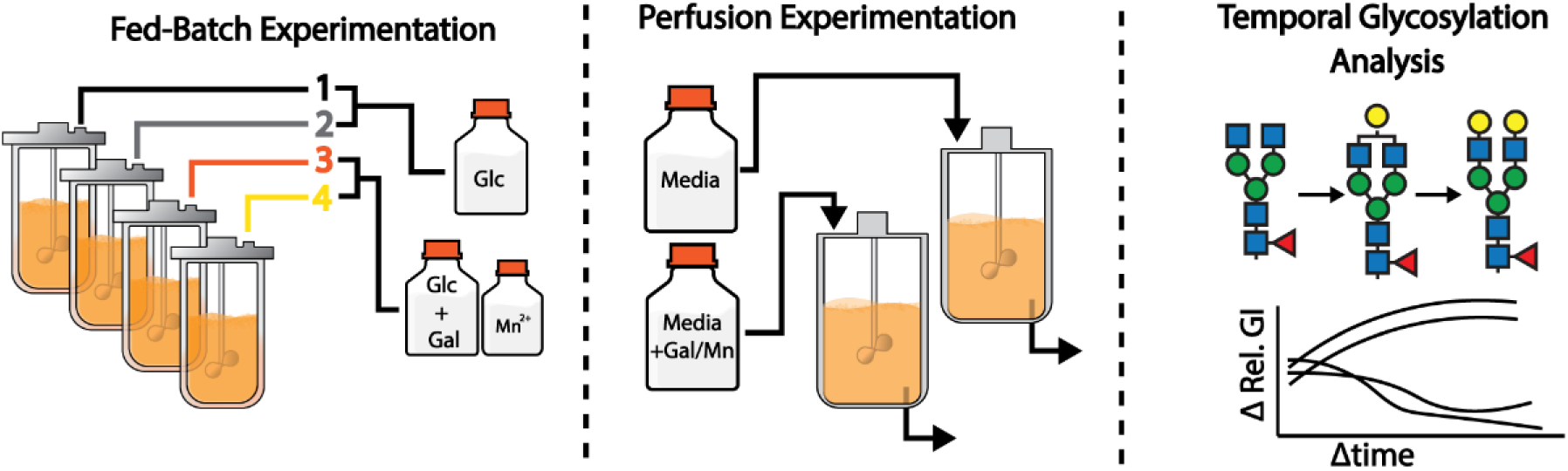
Study overview for fed-batch and perfusion-based cultures. For the fed-batch experiments, cultures were fed either daily (every 24 h) or on alternative days (every 48 h), and with either a bolus addition of glucose (control) or glucose and galactose (supplemented). For supplemented cultures, manganese was supplemented on the first feeding day to 1 μM. Perfusion-based experiments were perfused with either basal medium or basal medium spiked with galactose (1.8 g/L) and manganese (1 μM). A temporal analysis of mAb glycosylation and modulation of galactosylation will be done to understand the influence of feed and feeding regimen; values will be assessed in terms of the galactosylation index (GI) and separated glycoforms.

**Scheme 2:**
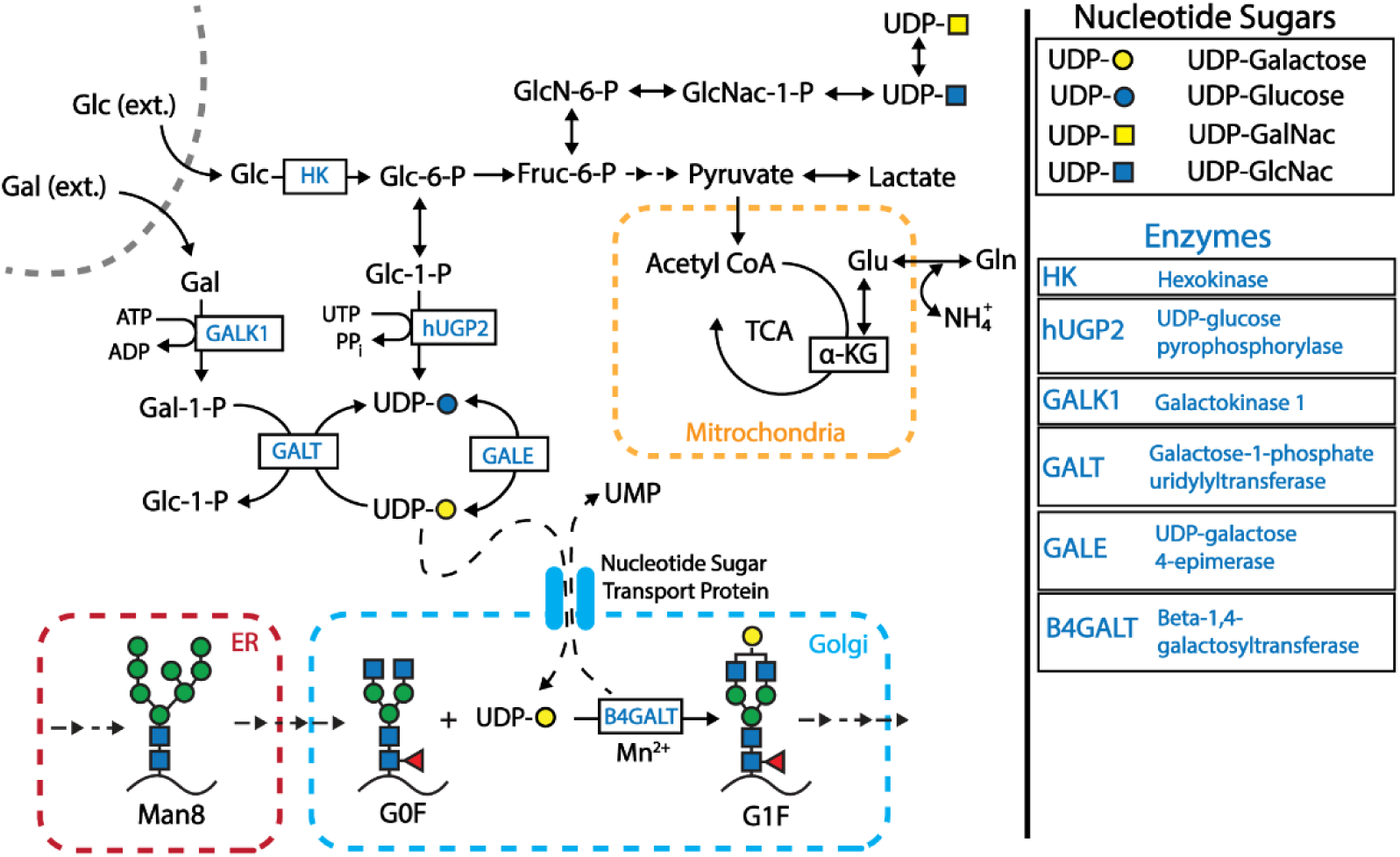
The interconnectivity of glucose and galactose metabolism. Scheme shows a brief depiction of the transport of glucose into glycolysis (top), the transport of galactose (middle) into the Leloir pathway, which branches to both glycolytic and glycosylation pathways. The bottom of the scheme depicts the mAb N-glycosylation pathway toward producing more mature glycoforms such as the G1F variant shown here.

## Graphical Abstract

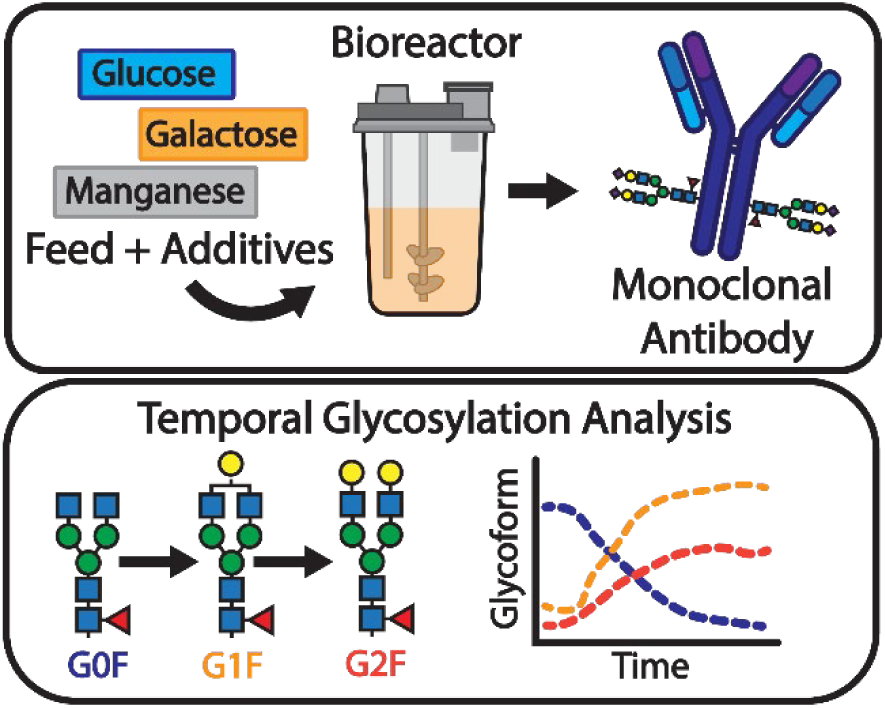

The study by Gyorgypal et al. investigates temporal effects of galactose and manganese on the glycosylation of Trastuzumab in fed-batch and perfusion bioprocessing modes, revealing the influence of feeding schedules on glycosylation patterns. The findings will aid in building process control schemes for mAb bioprocessing.

