## Supplemental Information for "Temporal Effects of Galactose and Manganese Supplementation on Monoclonal Antibody N-Linked Glycosylation in Fed-Batch and Perfusion Bioreactor Operation"

**Tables of Figures:**

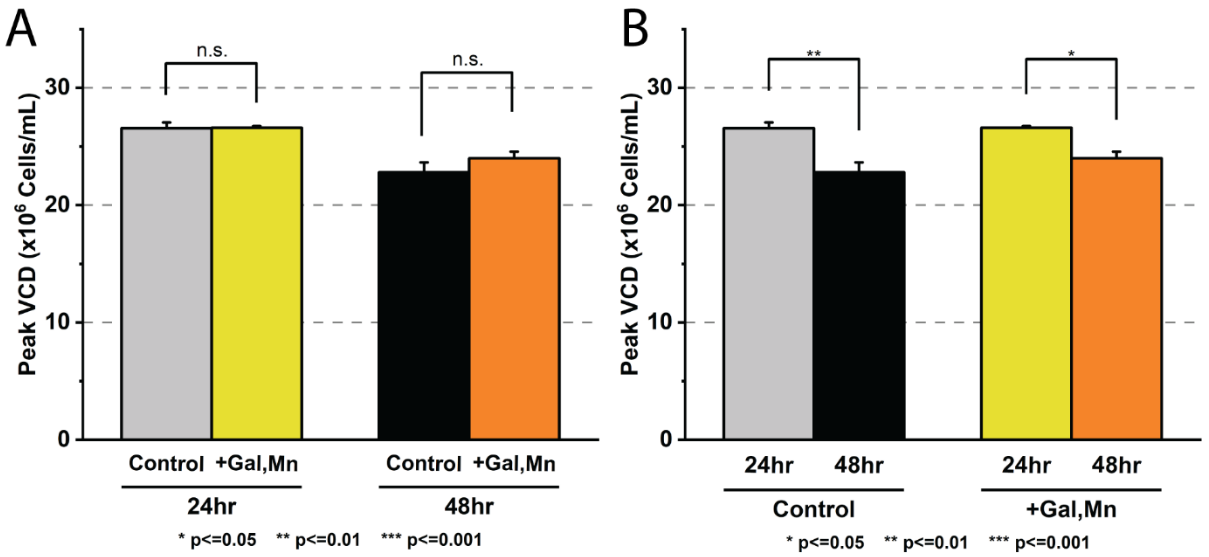

*Figure S1. Peak viable cell density comparison between fed-batch cultures.*

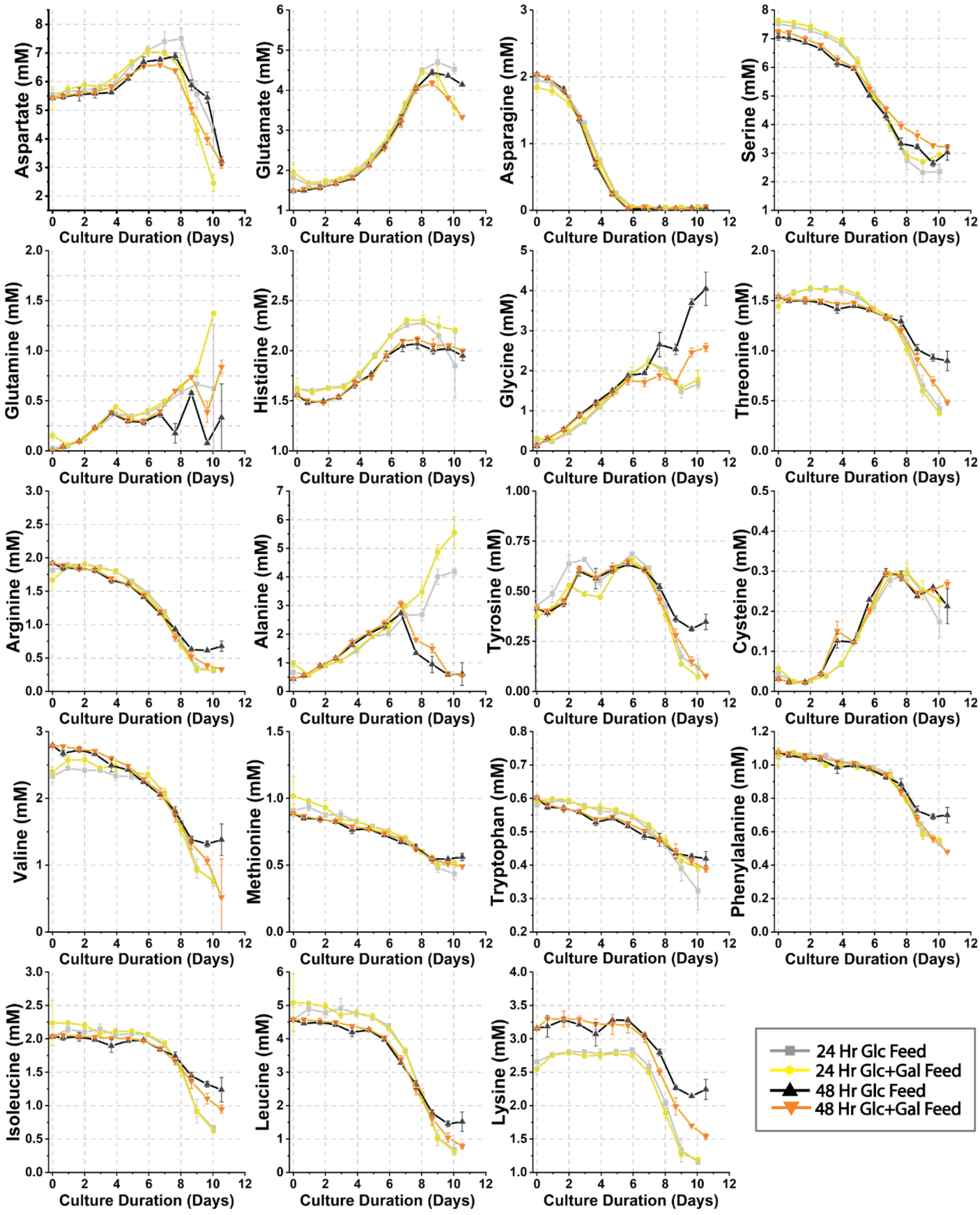

*Figure S2. Extracellular amino acid profile of fed-batch cultures.*

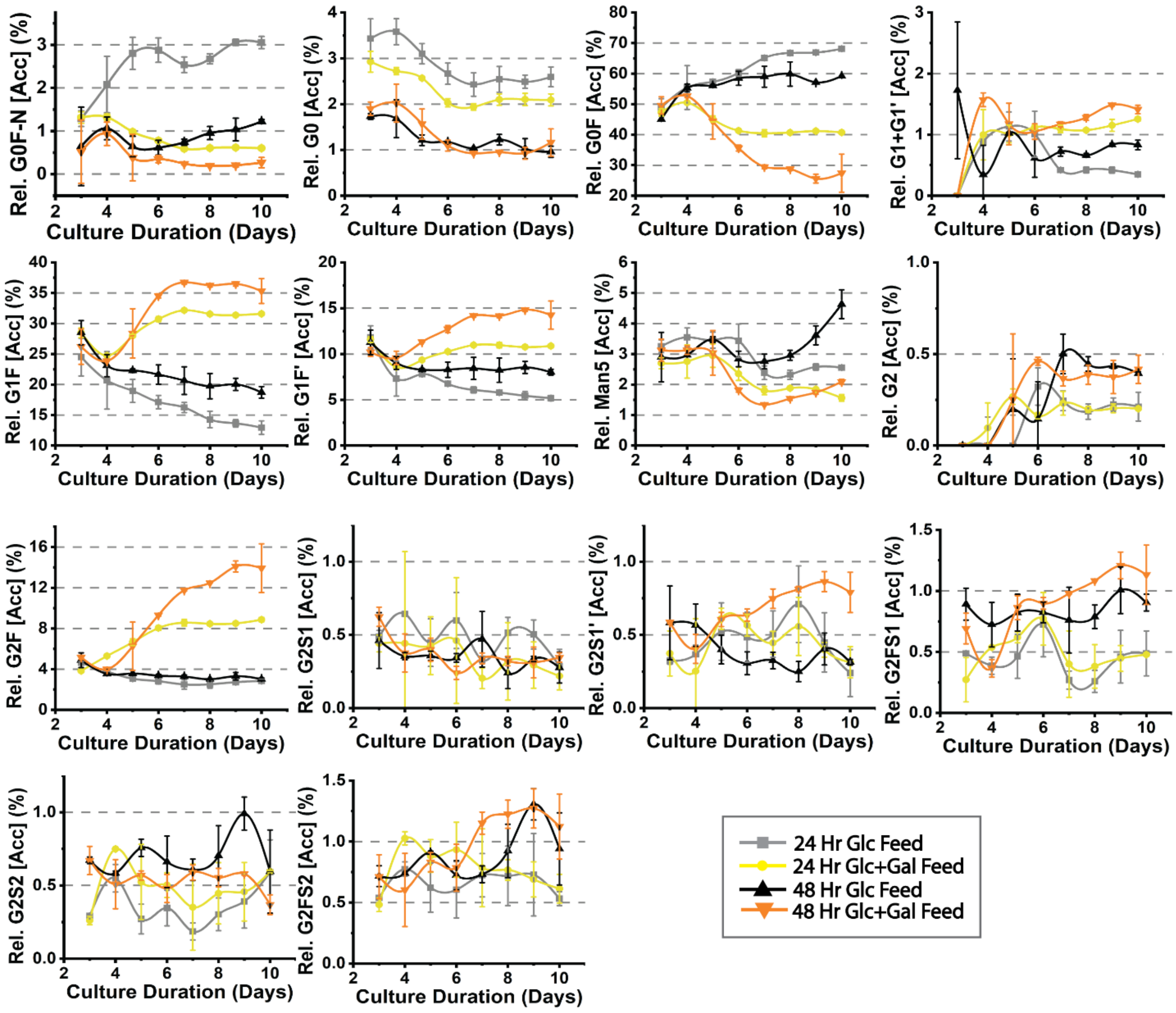

*Figure S3. Glycosylation of fed-batch cultures separated by glycoform.*

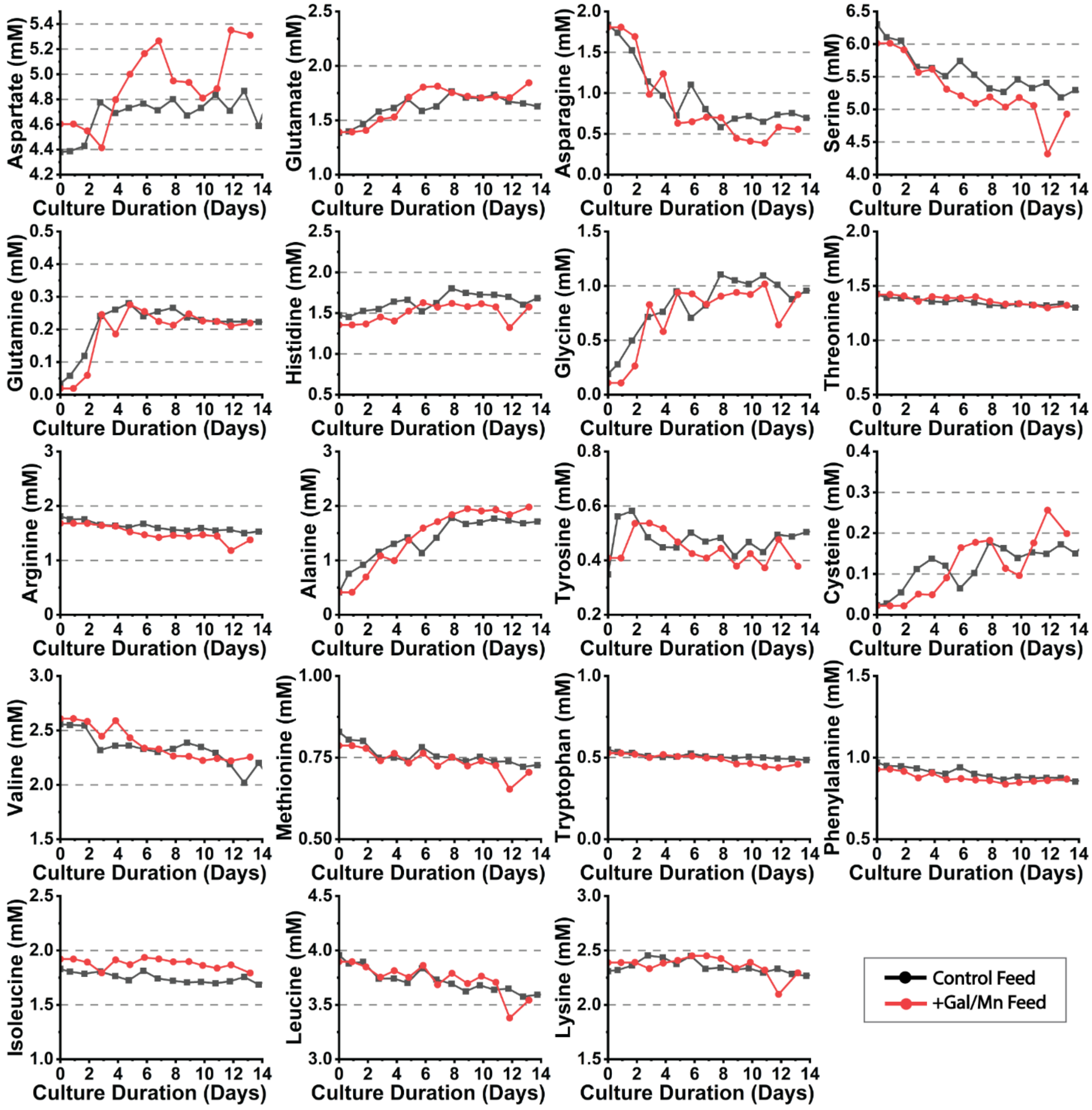

*Figure S4. Extracellular amino acid profile of perfusion cultures.*

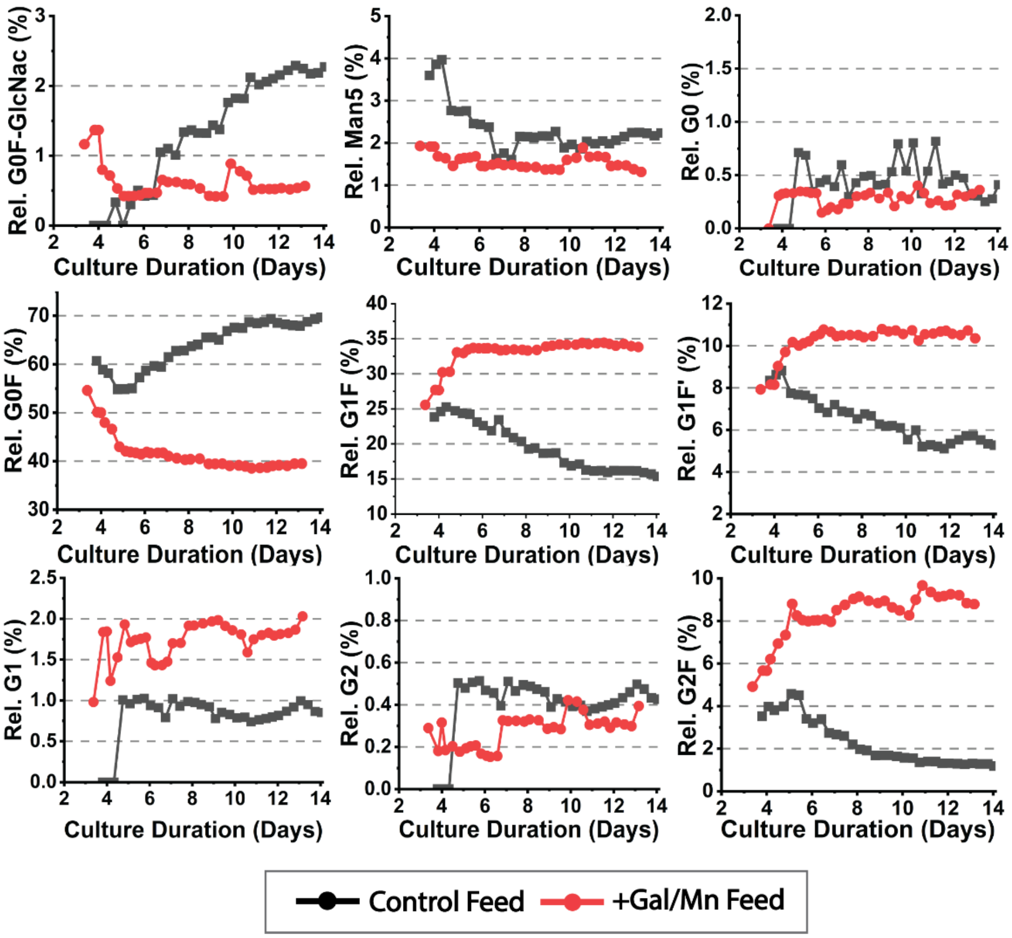

*Figure S5. Glycosylation of perfusion cultures separated by glycoform.*
